# Plasmotype condition nuclear pleiotropic effects on clock and fitness in barley

**DOI:** 10.1101/2021.11.02.466976

**Authors:** Eyal Bdolach, Manas Ranjan Prusty, Lalit Dev Tiwari, Khalil Kashkush, Eyal Fridman

## Abstract

In plants, the role of chloroplasts and mitochondria (plasmotype) in controlling circadian clock plasticity and overall plant robustness has not been elucidated. In this study, we investigated the rhythmicity of chlorophyll fluorescence (Chl F) clock output, and fitness in the field at optimal and elevated temperatures, in three different barley populations. First, we examined a reciprocal DH population between two wild barley (*Hordeum vulgare* ssp. *spontaneum*), in which we identified two pleiotropic QTLs (*frp2.1* and *amp7.1*) that modulate clock and fitness including conditioning of these effects by plasmotype diversity. In the second population, a complete diallel consisting of 11 genotypes (reciprocal hybrids differing in plasmotype), we observed a gradual reduction in plasmotype, ranging from 26% and 15% for Chl F and clock measurements to 5.3% and 3.7% for growth and reproductive traits, respectively. The third population studied was a collection of cytolines in which nine different wild plasmotypes replaced the cultivated Noga (*H. vulgare*) plasmotype. Here, the order and magnitude of the effects of the plasmotypes differed from what we observed in the diallel population, with the greatest effect of plasmotype diversity observed for clock period and amplitude. Comparison of the chloroplast sequences suggests several candidate genes in the plastid-encoded RNA polymerase (PEP) complex that may be responsible for the observed plasmotype effects. Overall, our results unravel previously unknown cytonuclear epistatic interactions that controls clock performance while also having pleiotropic effects on a plant field characteristics.

## INTRODUCTION

Plants are composed of cells in which three different organelle genomes co-evolved to cope with a dynamic environment: the genomes in nuclei, chloroplast and mitochondria (plasmotype). Phenotypic constraints promote selection of causal mutations in those three genomes and at the same time, interactions between genome products may impose epistatic relationship and co-evolution of adaptive gene complexes. In recent years, several studies have shown that phenotypic effects are related to the genetic diversity of the plasmotype and its interactions with the nucleotype (Joseph et al, 2013; Roux et al, 2016; Tang et al, 2014). An elegant use of haploid-inducer line available in *Arabidopsis* (*GFP-tailswap*) (Ravi et al., 2014) allowed generating a set of reciprocal and isogenic cybrids using several accessions, which was followed by phenotyping of metabolism and photosynthesis under different light conditions (Flood et al., 2020). Genetic analysis revealed that the nucleotype, plasmotype and their interaction accounted for 91.9, 2.9 and 5.2% of genetic variation, respectively, thus highlighting the importance of interactions between genomes. Moreover, variation explained due to cytonuclear epistasis was even higher (17.8%) for *Φ*_NPQ_ and changed significantly between different light regimes.

In crop plants, few reports exist on the contribution of cytonuclear interactions (CNI) to a plant’s phenotype, and even less to its effects on the plant’s phenotypic plasticity. Especially in grasses, the contribution of the plasmotype to yield and grain quality has been demonstrated (Frei et al., 2003; Sanetomo and Gebhardt, 2015). In cucumber, Gordon and Staub (2011) used reciprocal backcrosses between chilling-sensitive and chilling-tolerant lines to show that tolerance to reduced temperature is maternally inherited. Likely these traits are the result of a local adaption of the original wild alleles, since for example in bread wheat (*Trictium aestivum*) cytoplasmic influence on fruit quality is affected by genotype-by-environment interactions (Ekiz et al., 1998). Nevertheless, many of these examinations of alloplasmic lines, which contained cytoplasm from distantly related wild relatives showed that effects on agronomic traits (rather than protein quality) are not frequent (Frei et al., 2010). In maize, although cytoplasmic effects were not significant between the direct and reciprocal populations, the interactions among the cytoplasm and the nuclear quantitative trait loci (QTL) were detected for both days to tassel, and days to pollen shed (Tang et al., 2014), further enforcing the increased variation explained in *Arabidopsis* cybrids when cytonuclear interactions are included (Flood et al., 2020).

Circadian clock rhythms in plants are interwind with chloroplastic activities including photosynthetic phenotypes such as NPQ and ΦPSII that are important for plant productivity (Kromdijk et al., 2016). This connection led to the development of several high-throughput methods that measure the rhythmicity of the leaf chlorophyll fluorescence as a proxy to the period, phase and amplitude of the clock (Gould et al., 2009; Tindall et al., 2015; Dakhiya et al., 2017;). The ability to measure hundreds of plants allowed comparisons between species (Rees et al., 2019), identification of correlation for period and amplitude with temperature and soil composition (Dakhiya et al., 2017), as well as associating between naturally occurring circadian rhythm variation with clock gene loci in Swedish *Arabidopsis* accessions (Rees et al., 2021). Using the SensyPAM platform, which allows to infer clock output rhythmicity based on photosynthetic parameters (Bdolach et al., 2019), we recently analyzed wild, landraces, cultivars and interspecific barley populations. We showed that some of the nuclear loci that control the circadian rhythms were under selection during domestication, which could explain how modern crops lost the thermal plasticity of their clock (Prusty et al., 2021). Furthermore, pleiotropic effects of these drivers of clock (DOC) loci on grain yield under stress indicate the adaptive value of clock plasticity although the molecular, developmental, and physiological basis of this pleiotropy requires more experiments. Moreover, this study did not consider the possible role of cytoplasm diversity in manifesting these clock and pleiotropic effects on growth and reproductive fitness traits.

Here we follow up on the clock analysis of a reciprocal bi-parental doubled haploid (DH) population divided between genotypes carrying different plasmotypes from the Barley1K collection (from Ashkelon or Mount Hermon) (Hubner et al., 2009), and segregating for their nuclear genomes as well. We previously showed a significant difference of 2.2 h in the clock plasticity (delta of period) between the carriers of the different plasmotypes (Bdolach et al., 2019). In addition, we identified several nucleotype QTL that affected the period or the amplitude of the rhythmicity, based on ΦNPQlss measurements. In the current study, we intend to 1) extend the analysis of the plasmotype effects on fitness traits and test if there is pleiotropy between clock and life history traits, and 2) to extend the breadth of plasmotype diversity tested by adding additional crosses and more chloroplast sequencing information, and finally 3) to examine the potential of wild plasmotype diversity for modern crop breeding under optimal and high temperature.

## RESULTS

We wished to examine the effects of plasmotype diversity on growth and productivity of barley grown under ambient vs high temperatures and test possible relationship between circadian clock and growth plasticity. Previously, we described the generation of the ASHER doubled haploid population from two reciprocal hybrids between **Ash**kelon (B1K-09-07) and **Her**mon (B1K-50-04) wild barleys (Bdolach et al., 2019) This population of 121 genotypes is composed of 40 and 81 carriers of the B1K-09-07 and B1K-50-04 cytoplasms, respectively, whereby significant differences between two groups could be associated with plasmotype (mitochondria and chloroplast) variation. In addition to the homozygous ASHER population we developed an additional population by carrying out a set of reciprocal crosses between 11 wild barley accessions to achieve a full-diallel with few genotypes missing (see Methods). The rationale behind the diallel population is that any difference between pairs of hybrids can be associated with plasmotype differences between homozygous parental lines. Finally, we wanted to investigate the potential utility of the wild plasmotype for cultivated material and therefore generated and tested cytolines in the cultivar Noga background.

### Life history phenotypic responses in barley growing under high temperature

The plants of both the ASHER and diallel populations were phenotyped for life history traits and tested for differences between ambient temperature (AT) and high temperature (HT) from beginning of tillering stage until grain filling. We found that most of the life history traits were significantly different between the two environments in both populations (Fig. 1a and 1b). In ASHER (Fig 1a), Days to flowering (DTF) was significantly lower under HT (69 ±4.3 days) as compared to AT (100 ±3.6 days) (Fig 1a). The reproductive traits were higher under AT vs HT conditions (avg spike dry weight (ASDW)=0.89 ±0.13 vs. 0.7 ±0.12 gr and Spikes dry weight (SpDW)=7.8 ±1.9 vs 6.24 ±1.7 gr, respectively). Plant height (PH) at harvest was also higher under AT (124.9 ±8.5 cm) than under HT (108.1 ±7.8 cm) although vegetative dry weight (VDW) was lower under AT, i.e. 13.1 ±3.5 gr vs 14.4 ±4.7 gr in HT. As a result, Total dry matter (TDM) was not significantly different between environments (AT, 20.8 ±5.6 and HT, 20.5 ±6 gr). Spike length (SL) also was not significantly different between environments (AT, 9.7 ±0.9 and HT, 9.7 ±1 cm) while the variation in the number of spikes per plant (calculated as the coefficient of variation (CV), SLCV) is significantly lower (more stable) under AT (AT, 8.3 ±1.6 and HT, 11.03 ±4.2 %). This could also be viewed based on the wider distribution in the SLCV under HT. It is interesting to note that while the HT affected the SpDW and included in fact reproductive output loss, the VDW worked in an opposite manner including gain in the weight of the non-reproductive parts under HT.

**Figure 1:**
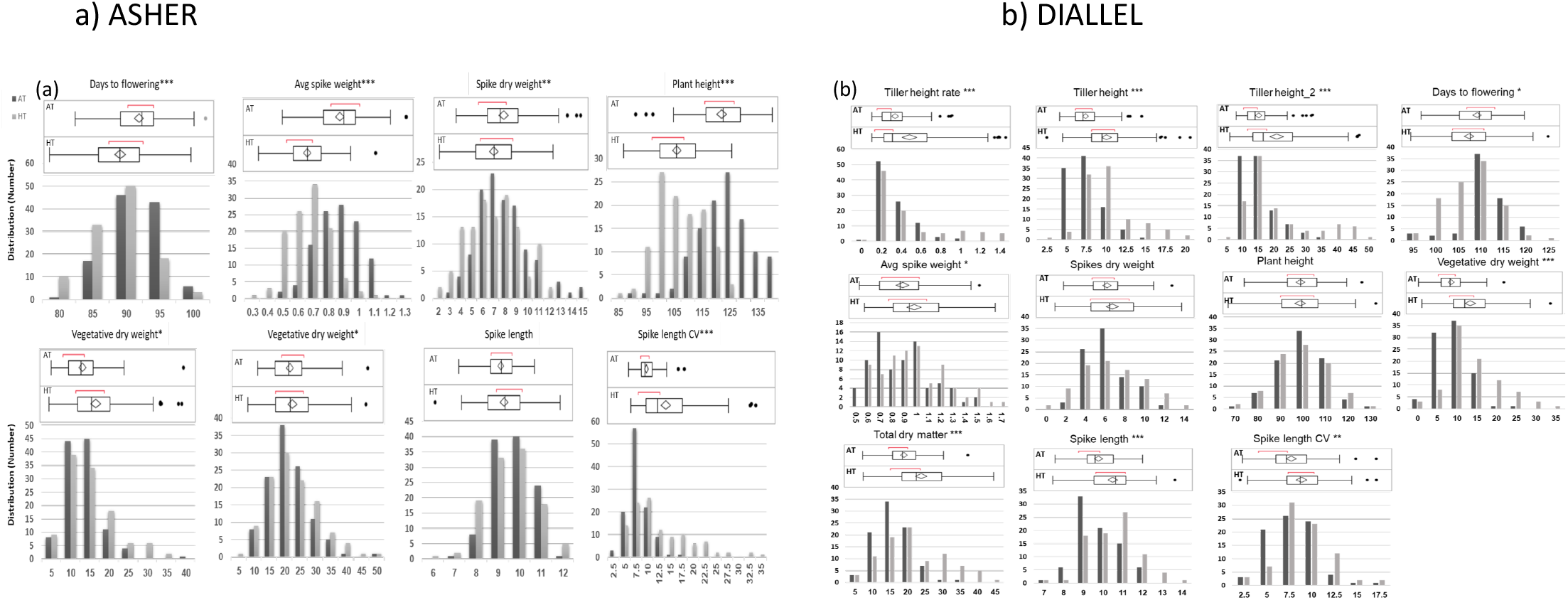
Mild increase in temperature has significant effect on plant performance in the field. Distribution and box plot of life history traits under ambient temperature (AT, black) and high temperature (HT, gray) (see Fig. S1 for differences between AT and HT) for **a)** ASHER DH population and **b)** full-diallel. Life history traits include Tiller height at two time points and the rate between them, Average Spike dry weight, Days to flowering (DTF), Spike length (SL), Spikes dry weight (SpDW), Plant height (PH), Vegetative dry weight (VDW) and Total dry matter (TDM). For each student’s t-test between HT and AT, the p value is depicted as *: P<0.05, **: P<0.01 or ***: P<0.001

Unlike in the ASHER population, in the diallel experiment we noticed almost identical DTF between HT and AT (DTF=-109.9 ±5.6 and 111.73 ±4.97 days, respectively) (Fig 1b). Similarly, unlike the significant effects of the thermal environment on the vegetative traits, the reproductive traits were less affected. ASDW is significantly yet mildly lower under AT (0.93 ±0.24 gr) than under HT (1.03 ±0.27 gr). For SpDW we could not detect a significant difference between environments (7.13 ±2.2 and 7.35 ±3.16 gr, respectively). PH is also not significantly different between environments (104.06 ±10.17 cm in AT and 104.04 ±12.05 cm in HT), as compared to VDW and TDM that were significantly lower under AT (11.29 ±4 and 18.5 ±5.5 gr) than HT (16.38 ±6.8 and 24.13 ±9.08 gr). SL and SLCV are significantly lower under AT.

To summarize, in both field experiments of the two populations (ASHER and diallel) the plants on average accumulated lesser VDW and showed higher stability between spikes, i.e. lower VDW and SLCV, under AT. In addition, doubled haploid plants were flowering earlier under HT.

### Plasmotype effects on plasticity of life history traits, circadian clock, and Chl F

The ASHER population is composed of two sub populations, each carrying either the Ashkelon or Hermon plasmotypes (Bdolach et al., 2019). On average, carriers of the Hermon plasmotype flowered significantly earlier than the Ashkelon types under both HT and AT however, in both subpopulations, this manifested in the same acceleration of flowering by more than 2 days (Fig. 2a). For SL there is no difference (Fig 2b), but for the SLCV Hermon plasmotype is linked with lesser stability (Fridman, 2015), i.e. higher CV under HT (14.45% under HT vs 8.8% under AT) as compared to Ashkelon (12.33% under HT vs 9.32% under AT) (Fig. 2c). Perhaps the most interesting comparison is between the total vegetative and reproductive outputs. The carriers of the Ashkelon plasmotype are on average very plastic for the plant biomass and for the derived total dry matter (Fig. 2d and 2e), while the Hermon plasmotype types are relatively stable for the biomass and respond significantly with reduction of the spikes dry weight under heat (Fig. 2f). This is in comparison to the relative stable SpDW of the Ashkelon types (Fig. 2f).

**Figure 2:**
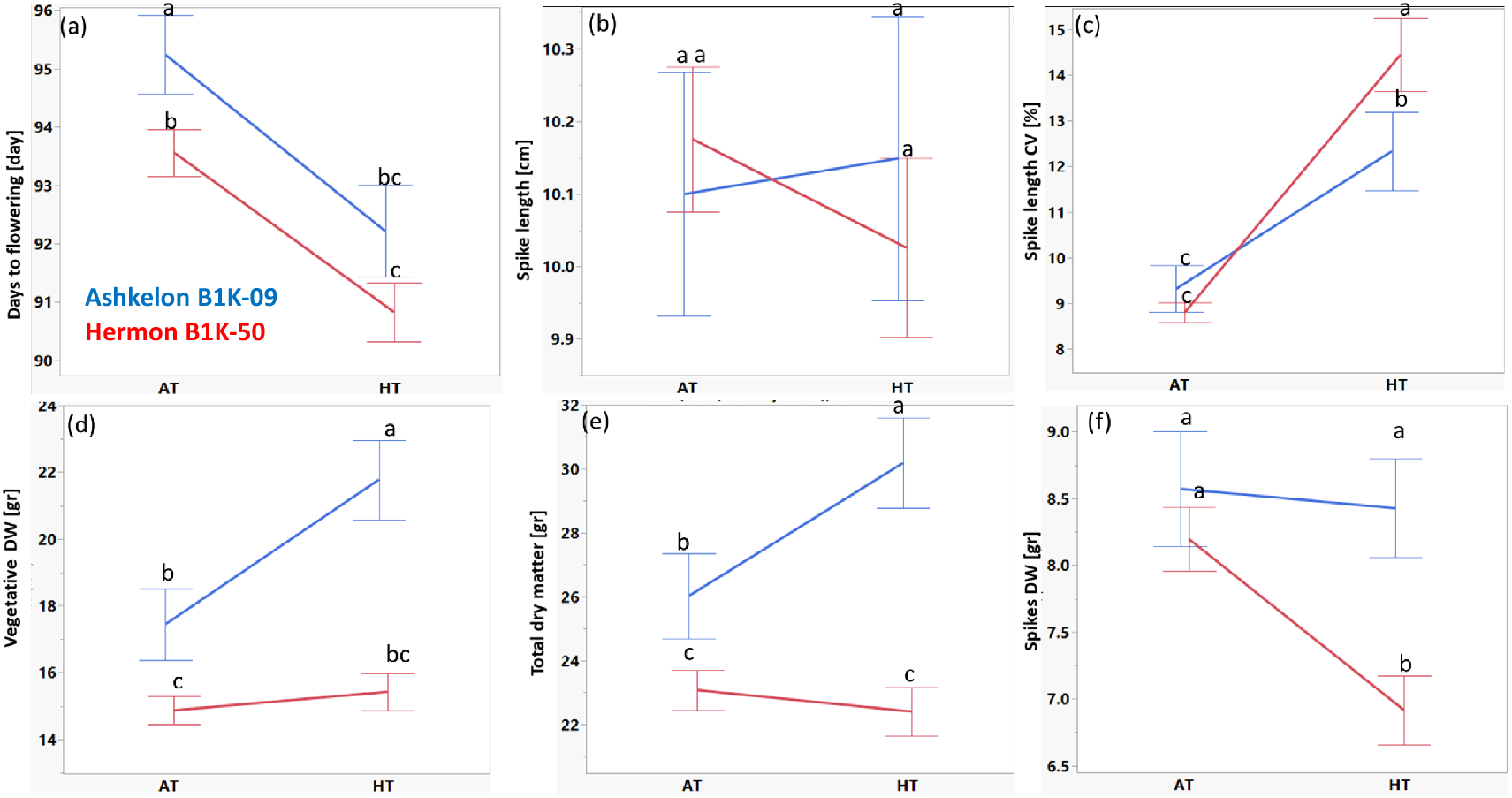
Thermal plasticity of life history traits is under cytoplasmic control in wild barley. Reaction norms of life history traits depicting the average responses of the two parental plasmotypes to mild heat. Differential response between the carriers of Ashkelon (blue) and Hermon (red) plasmotype for **a)** Days to flowering (DTF), **b)** Spike length (SL), **c)** Spike length CV (SLCV), **d)** Vegetative dry weight (VDW), **e)** Total dry matter (TDM) and **f)** Spikes dry weight (SpDW). Levels not connected by same letter are significantly different in student’s t-test (P<0.05).

The reciprocal nature of the hybrids in the full-diallel allowed us to group the F1 plants into different plasmotype subpopulations and different male parent subpopulations (representing the nucleotype). One-way ANOVA for each of these two sub-populations indicated larger percentage variation explained (PVE) by the nucleotype (male donors), in comparison to differences between plasmotype (female donors) for few traits (Table 1). For example, for the ASDW under HT (PVE=41% for nucleotype vs PVE=27% for plasmotype) and DTF under AT (PVE=43% vs PVE=32%) and HT (PVE=37% vs 28%). For majority of the life history traits we found higher variation that explained by the plasmotype than nucleotype under both temperatures (AT and HT): for PH, PVE=39% vs 30% and 33% vs 21% under AT and HT, respectively. This was true also for reproductive output, e.g SpDW which showed higher variance between plasmotypes in AT (PVE=35% vs 21%) and to a lesser extent under HT (PVE=23% vs 19% between plasmotype and nucleotype contributions).

**Table 1:** One-way ANOVA for plasmotype vs nucleotype effects in the reciprocal diallel population. Tiller height at two time points and the rate between them, Days to flowering (DTF), Average Spike dry weight (ASDW), Plant height (PH), Spike length (SL), spike length CV, Spikes dry weight (SpDW), Total dry matter (TDM) and Vegetative dry weight (VDW) in the nethouse under ambient temperature (AT) and high temperature (HT). For clock and Chl F traits: Amplitude, Period, delta Amplitude (dAMP), delta Period (dPeriod), Fv/Fm, F/Fmlss, NPQlss and Rfd in SensyPAN under optimal temperature of 22°C (OT) or high temperature of 32°C (HT) and the delta HT-OT.

| Trait | Type | male<br>parent<br>Prob > F | male parent<br>PVE [%] | Plasmotype<br>Prob > F | Plasmotype<br>PVE [%] |
| --- | --- | --- | --- | --- | --- |
| TH_1_AT | Growth | 0.0068 | 24 % | <.0001 | 41 % |
| TH_1_HT | Growth | <.0001 | 37 % | <.0001 | 36 % |
| TH_2_AT | Growth | <.0001 | 34 % | <.0001 | 38 % |
| TH_2_HT | Growth | <.0001 | 36 % | <.0001 | 46 % |
| TH rate_AT | Growth | <.0001 | 34 % | 0.0002 | 32 % |
| TH rate_HT | Growth | <.0001 | 33 % | <.0001 | 47 % |
| ASDW_AT | Reproductive | <.0001 | 43 % | <.0001 | 35 % |
| ASDW_HT | Reproductive | <.0001 | 41 % | 0.0037 | 27 % |
| DTF_AT | Reproductive | <.0001 | 43 % | 0.0002 | 32 % |
| DTF_HT | Reproductive | <.0001 | 37 % | 0.0008 | 28 % |
| PH_AT | Growth | 0.0007 | 30 % | <.0001 | 39 % |
| PH_HT | Growth | 0.0266 | 21 | 0.0001 | 33 % |
| SL_AT | Reproductive | 0.0006 | 31 % | <.0001 | 42 % |
| SL_HT | Reproductive | 0.0111 | 23 % | 0.0007 | 30 % |
| SLCV_AT | Reproductive | 0.0215 | 22 % | 0.0912 | 18 % |
| SLCV_HT | Reproductive | 0.2076 | 15 % | 0.1434 | 16 % |
| TDM_AT | Reproductive | 0.0879 | 18 % | <.0001 | 35 % |
| TDM_HT | Reproductive | 0.3174 | 13 % | 0.0007 | 29 % |
| VDW_AT | Growth | 0.1108 | 17 % | <.0001 | 35 % |
| VDW_HT | Growth | 0.1965 | 15 % | 0.0003 | 31 % |
| SpDW_AT | Reproductive | 0.028 | 21 % | <.0001 | 35 % |
| SpDW_HT | Reproductive | 0.0566 | 19 % | 0.0094 | 23 % |
| Amplitude_OT | Clock | 0.1819 | 24 % | 0.1325 | 26 % |
| Amplitude_HT | Clock | 0.2046 | 23 % | 0.136 | 26 % |
| Period_OT | Clock | 0.2694 | 22 % | 0.387 | 19 % |
| Period_HT | Clock | 0.2715 | 22 % | 0.0184 | 34 % |
| dAMP | Clock | 0.1892 | 24 % | 0.0369 | 32 % |
| dPeriod | Clock | 0.7274 | 13 % | 0.3768 | 20 % |
| Fv/Fm_OT | Chl F | 0.736 | 14 % | 0.2647 | 23 % |
| Fv/Fm_HT | Chl F | 0.4621 | 19 % | 0.3934 | 20 % |
| Fv/Fmlss_OT | Chl F | 0.9231 | 09 % | 0.0834 | 29 % |
| Fv/Fmlss_HT | Chl F | 0.8984 | 10 % | 0.2307 | 24 % |
| NPQlss_OT | Chl F | 0.4743 | 18 % | 0.0386 | 32 % |
| NPQlss_HT | Chl F | 0.1641 | 26 % | 0.0551 | 31 % |
| Rfd_OT | Chl F | 0.1015 | 28 % | 0.1697 | 25 % |
| Rfd_HT | Chl F | 0.0414 | 32 % | 0.0327 | 33 % |

We also included similar clock analysis to the hybrids of the diallel as we previously conducted for the ASHER population, i.e. under optimal (OT) and high temperatures (HT) environments using SensPAM (See Methods; Bdolach et al., 2019). The clock rhythmicity (amplitude and period) is based on NPQlss measurements for three days under constant light (Dakhiya et al., 2017) (. The clock amplitude was significantly higher under HT (0.03 ±0.01) compared to OT (0.015 ±0.006) (Fig 3a). Regarding the clock period, we observed significantly higher values under OT (24.9 ±2.6 h) in comparison to HT (23.3 ±1.9 h; Fig 3b). This clock plasticity is similar to the one described for the ASHER population (Bdolach et al., 2019) with acceleration of the rhythmicity under higher temperatures. Fv/Fm is significantly higher under HT (0.93 ±0.01) in comparison to OT (0.92 ±0.01; Fig 3c) and significantly different for Fv/Fmlss (0.9 ±0.01 in OT vs 0.91 ±0.01 in HT; Fig 3d). NPQlss and Rfd are significantly different under OT in comparison to HT (NPQlss 0.66 ±0.1 vs 0.43 ±0.08 and Rfd 1.6 ±0.2 vs 1.18 ±0.18; Fig 3e and f). Overall, these F results suggest that under HT. photosynthesis is more efficient.

**Figure 3:**
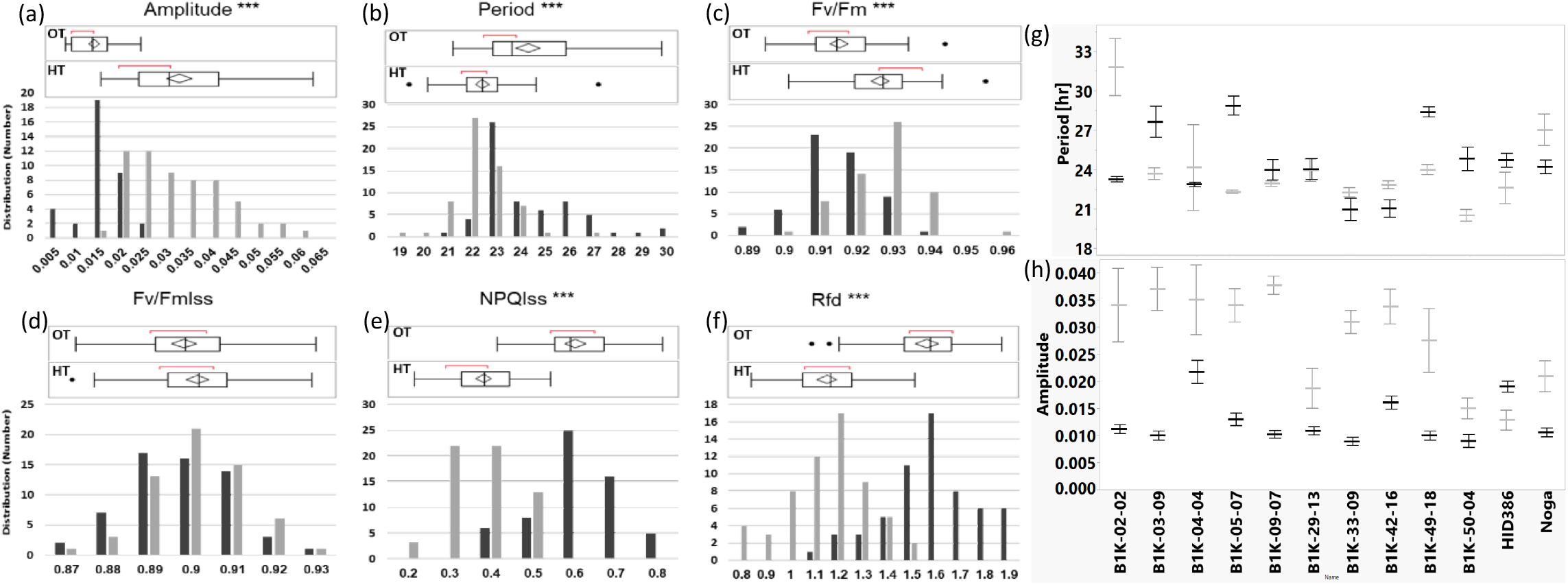
Chlorophyll florescence and circadian clock rhythmicity in barley diallel grown under OT and HT. Distribution and box plot of clock output rhythmicity: **a)** Amplitude, and **b)** Period, and for mean chlorophyll florescence traits:, **c)** Fv/Fm, **(d)** Fv/Fmlss, **e)** NPQlss and **f)** Rfd under optimal temperature (OT, black) and high temperature (HT, gray) in the reciprocal diallel population. For each student’s t-test, the p value is depicted as *: P<0.05, **: P<0.01 or ***: P<0.001. The means of clock **g)** period and **h)** amplitude under optimal temperature (OT, black) and high temperature (HT, gray) in SensyPAM for the *H. spontaneum* accessions parental accessions of the diallel: B1K-02-02 (Yerucham), B1K-03-09 (Michmoret), B1K-04-04 (Ein Prat), B1K-05-07 (Neomi), B1K-09-07 (Ashqelon), B1K-29-13 (Mount Arbel), B1K-33-09 (Mount Harif), B1K-42-16 (Jordan Canal), B1K-49-19 (Mount Eitan), B1K-50-04 (Mount Hermon), HID386 (Kisalon, Israel) and the cultivar Noga.

Differences between the contributions of plasmotype and nucleotype to clock traits in the diallel are also found (Table 1), but to a lesser extent than for fitness traits. This includes higher PVE by the plasmotype of 34% compared to 22% by nucleotype for period under HT. Similarly, the delta amplitude variation between hybrids is better explained by plasmotype (32%) compared to nucleotype (24%) (Table 1).

### Relationship between plasmotype and nuclear diversity and pleiotropic effects on circadian clock and life history traits

In the diallel population we tested if there are reciprocal hybrids that significantly differ for life history and SensyPAM traits (Fig 4). We clustered phenotype measured in the nethouse as growth or reproductive ones and traits measured with SensyPAM as clock or Chl F parameters. The percentage of differing pairs of reciprocal hybrids is the highest for Fv/Fm under OT (44.8%) and the lowest is zero hybrids for the difference of DTF under AT. If comparing the traits according to our clustering, we could see that the mean number of differing reciprocal hybrids is highest for Chl F (26.3%), and second for clock traits (15%), while growth and reproductive traits are falling behind with 5.23% and 3.73%, respectively.

**Figure 4:**
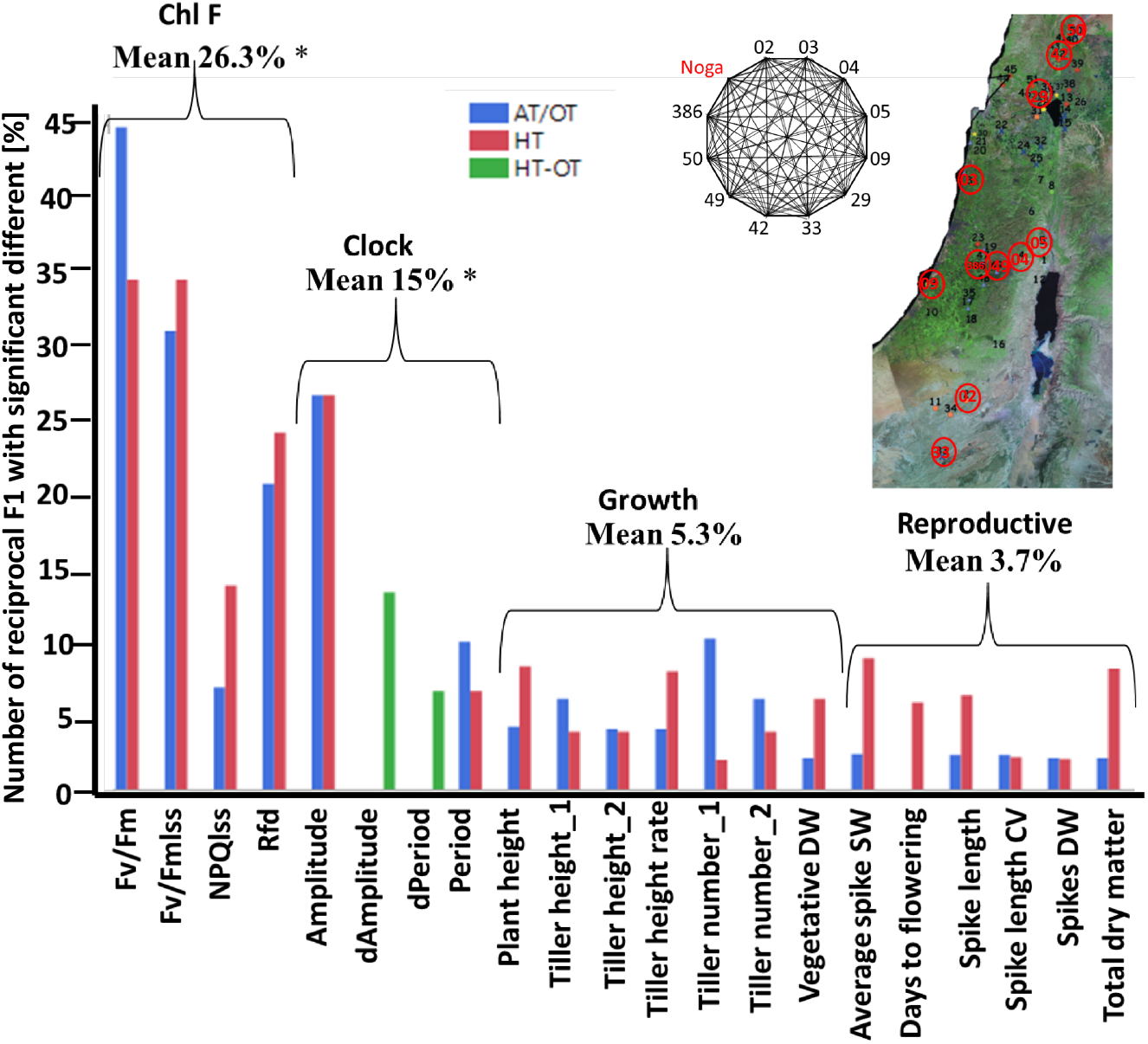
Proportion of crosses with significant difference between reciprocal hybrids for phenotypic traits, under OT or AT and HT. Life history traits include: Tiller height (TH) and number (TN) at two time points and the rate between them, Days to flowering (DTF), Average Spike dry weight (ASDW), Plant height (PH), Spike length (SL), spike length CV (SLCV), Spikes dry weight (SpDW), Total dry matter (TDM) and Vegetative dry weight (VDW) for plants in the nethouse under ambient temperature (AT) and high temperature (HT). For clock and Chl F traits: Amplitude, Period, delta Amplitude (dAMP), delta Period (dPeriod), Fv/Fm, F/Fmlss, NPQlss and Rfd in SensyPAM under optimal temperature of 22°C (OT) or high temperature of 32°C (HT) and the delta HT-OT. The mean for each type of traits is depicted: Growth or reproductive traits in the greenhouse experiment, and clock or Chl F in the SensyPAM.

One direct attempt to test for the relationship between circadian clock behavior under OT and HT to that of the plants fitness in the field is to perform simple linear regression between the different traits (Table S1). We found the highest positive clock-fitness correlations exist between period under HT and the two time points of tiller height measurements at the tillering stage and the rate between them (r= 0.432 and 0.535). Period under HT is also positively correlated to ASDW and SL under AT (r=0.46 and 0.42, respectively). This suggests that faster growth under both AT and HT is connected to longer clock periods under HT. Negative correlation was found between thermal responses of the amplitude (delta amplitude (dAMP) between HT and AT) and DTF under AT and HT (r= −0.44 and −0.5), i.e. the greater the value for dAMP, the earlier plants reach flowering. We also identified highly positive correlation between dAMP and SpDW under AT (r=0.45). Finally, we found negative correlations between Fv/Fm and Fv/Fmlss under both OT and HT and PH under AT and HT and a smaller positive correlation with NPQlss.

Another way to look into the relationship between clock and growth traits is through pleiotropy (same locus affecting several traits; (Chen and Lübberstedt, 2010)). We previously obtained circadian clock phenotypes and used this data for a genome scan in the ASHER DH population. This allowed us to identify several QTLs linked with variation in amplitude, period and heat responses (delta of the traits) and to verify them by segregation analysis (Bdolach et al. 2019). Here, we performed a similar genome scan analysis with the new set of field phenotypes (see Methods), and we also tested genetic models that include plasmotype and nuclear QTL interactions (cytonuclear, or GxG interactions). The genome scan identified several major QTL that are associated with the different traits, including few that we found to be pleiotropic (Fig 5a). On chromosome 2 we found several significant QTLs for SDW under HT (LOD=3.04), for VDW under HT and QxE (LOD= 8.86 and 8.68, respectively). In addition, we positioned a major QTL for PH under HT (LOD=6.35) which resides at the coordinates as the *frp2.1* locus that we linked previously with circadian clock period plasticity in the same population (Bdoalch et al., 2019). This major pleiotropic QTL resides between positions 698,875,542 and 702,308,910 (Morex v1). In the previous work, we identified this locus based on a threshold model, where we translated the period plasticity phenotype of the DH lines into a binary vector. For PH, there was no significant difference between carriers of the two alleles under AT, but under HT the Ashkelon allele was associated with a significant higher PH than the Hermon allele (114.65 cm vs 108.53 cm, respectively) (Fig 5b). In addition, carriers of the Ashkelon allele had on average a significantly higher VDW under HT vs. AT (24.57 gr and 19.93 gr, respectively), which is significantly higher than the results obtained for Hermon allele carriers under both treatments (14.86 gr in AT and 16.01 gr in HT). VDW also showed a significant QxE interaction explaining 50% of the trait variation (Fig. 5c). Another mild pleiotropic QTL is *amp7.1* individually found for Amplitude under HT and QxE on chromosome 7 (498,472,330-510,903,725; Morex v1). In this study we found this QTL for SL under HT (LOD= 8.47) (Fig 5a). For this QTL, SL of the Hermon allele carriers is higher in both treatments than those DH carrying the Ashkelon allele (10.4 and 10.34 cm vs 9.66 and 9.5, respectively) (Fig 5d).

**Figure 5:**
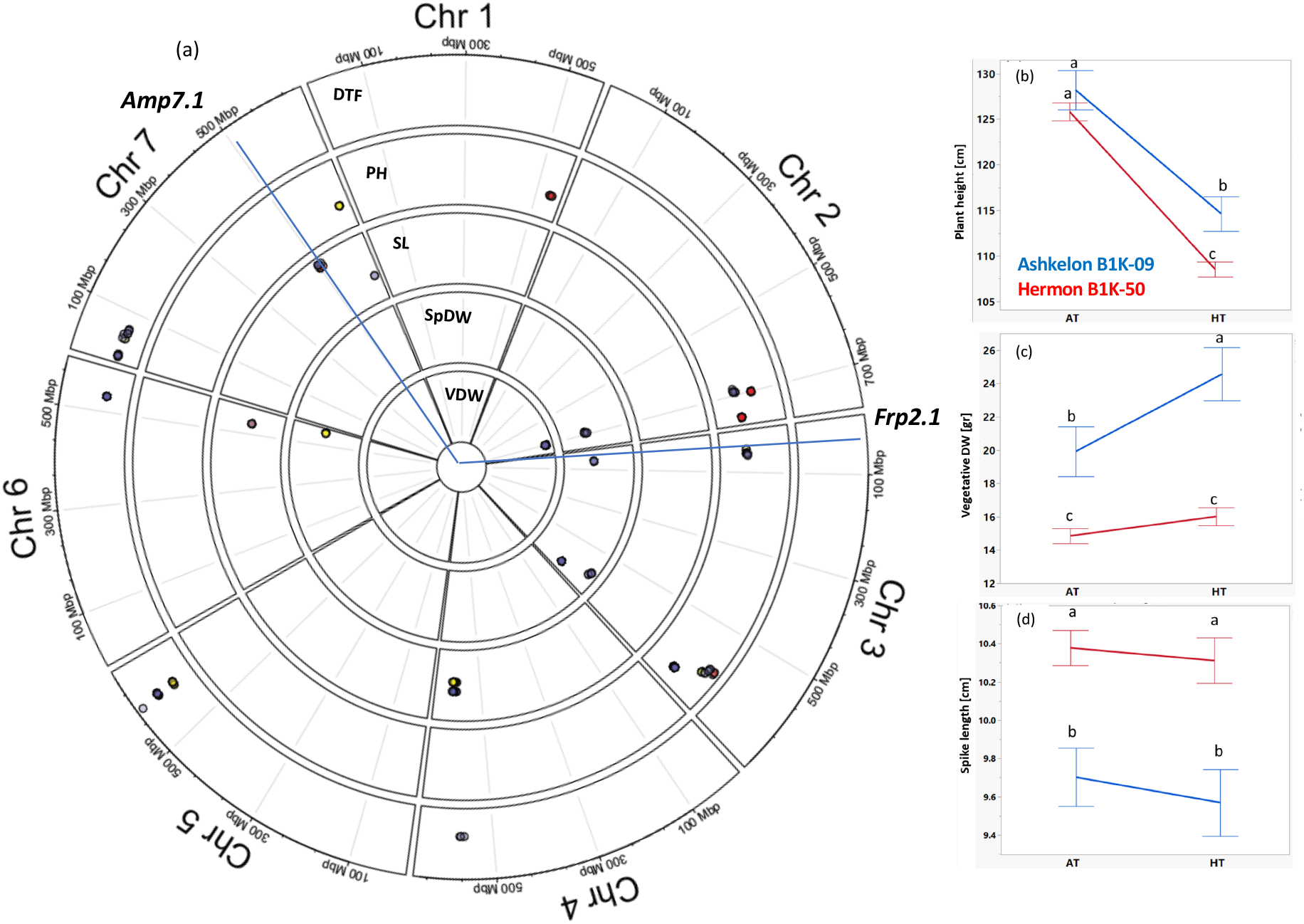
Pleiotropic nucleotype QTL underlying life history traits plasticity. Circos plot depicting loci with significant effects under AT, HT and loci showing GxE interaction with thermal environment. **a)** Circus of LOD in the ASHER DH population for life history traits. From outer to inner lane: Days to flowering (DTF), Plant height (PH), Spike length (SL), Spikes dry weight (SpDW) and Vegetative dry weight (VDW) under ambient temperature (AT-yellow), high temperature (HT-red) and GxE (blue). Reaction norms of *frp2.1* locus with pleiotropic effects on **b)** plant height (PH) and **c)** Vegetative dry weight (VDW), and of **d)** *amp7.1* for spike length. Red and blue lines depict mean values of lines homozygous for the Hermon or Ashkelon alleles, respectively. Levels not connected by same letter are significantly different in student’s t-test (P<0.05).

Finally, we tested the possible interactions between plasmotype and nuclear QTLs, i.e. whether the plasmotype diversity is conditioning the effects of the nuclear QTL. Under AT, for both VDW and SpDW the effect of the *frp2.1* QTL on Ashkelon vs. Hermon phenotypes was severely dependent on the DH plant carrying the Ashkelon plasmotype (Fig. 6a and 6c). This epistatic effect of the plasmotype over the nuclear locus was also found for VDW under HT (Fig. 6a) but did not appear when looking at the reproductive output under HT (Fig. 6d). Superimposing a reaction norm onto the interaction plot (Fig. 6b-d) indicates that recombination between the two loci (plasmotype and *frp2.1*) leads to an opposite behavior for the reproductive output. While no significant changes were observed between the slopes of the different plasmotype-*frp2.1* combinations for AT and HT, the carriers of the Hermon plasmotype with the Ashkelon allele at *frp2.1* showed increased reproductive output under HT compared to AT, in an opposite manner to the combination of Hermon-Hermon in both nuclear and cytoplasmatic loci (Fig. 6f).

**Figure 6:**
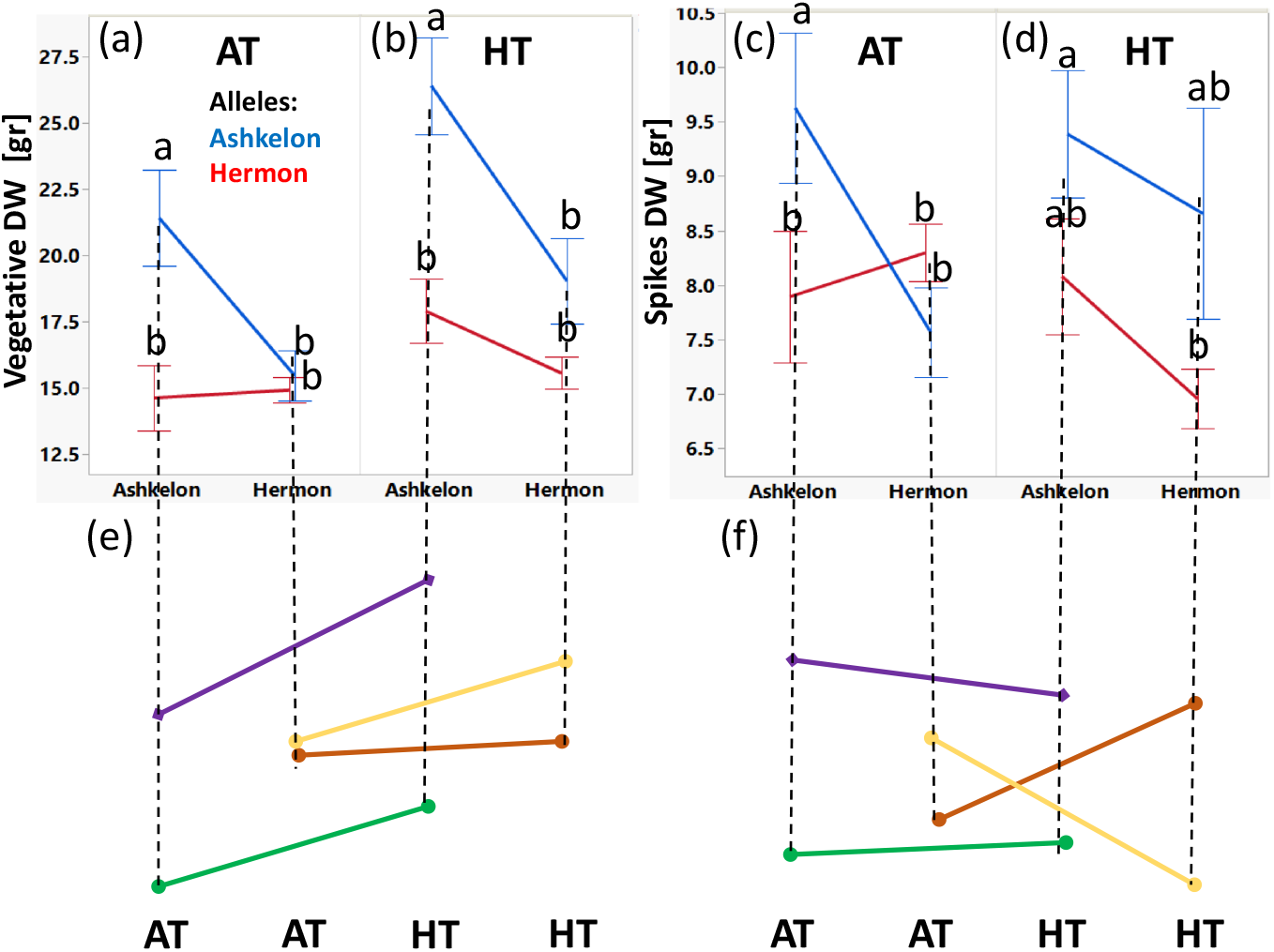
GxGxE interactions between plasmotype and *frp2.1* nuclear QTL under ambient and high temperatures in ASHER DH population. Cytoplasm by nuclear (GxG) QTL interaction plots under **a)** ambient (AT) and **b)** high (HT) temperatures for vegetative DW. Similarly, interaction plots under **c)** ambient (AT) and **d)** high (HT) temperatures for spikes DW. Plasmotype is depicted in x axis (Ashkelon or Hermon), and red or blue lines illustrate the Hermon or Ashkelon alleles in *frp2.1*. Reaction norms of the different Plasmotype-*frp2.1* locus-combinations between AT and HT for **e)** vegetative DW, and **f)** Spikes DW. Green, Ashkelon (Plasmotype)-Hermon (*frp2.1*); Purple, Ashkelon-Ashkelon; Brown, Hermon-Ashkelon; Orange, Hermon-Hermon.

Additional significant cytonuclear interactions for a pleiotropic QTL, on clock and growth, was found for *amp7.1* and its combined effects on days to flowering. In this *amp7.1*-plasmotype combination, we observed significant cross-over effects (Malosetti et al., 2013), i.e. changes in the order of the *amp7.1* genotypes between the two different plasmotypes (figure *2*). However, in this combination, the reaction norms looked identical between the four cytonuclear combinations.

### Plasmotype variation in Cytolines in clock rhythmicity, Chl F and life history traits

To test the hypothesis that the plasmotype has a role in regulating phenotypic diversity and could be utilized for breeding heat tolerance, we backcrossed several wild barley accessions with a cultivated elite line while keeping the wild plasmotype, in order to obtain nearly-isogenic cytolines (see Methods). The cytolines were tested in the SensyPAM under OT and HT for clock rhythmicity (measured from NPQlss) and Chl F, and in the nethouse for life history traits under AT and HT (Fig S1c). We calculated delta values between HT to OT or AT, in SensyPAM as well as in the nethouse, for each trait as a measure of the thermal plasticity (Fig. 7). We found significant differences between cytolines for clock period under HT (P=0.004) and not under OT (P=0.17) (Fig S3a). Notably, the single cytoline that decelerated the clock significantly between OT and HT is the one carrying the B1K-03 plasmotype (Noga^03^, decelerated by 4.3h, P<0.05; Fig 7a). Unlike the relative uniformity between cytolines for period under OT and HT (Fig. S3a) the variance between cytolines for clock amplitude was significant under both OT (P=0.0001) and HT (P=0.006) (Figure S3b). Most of the cytolines showed a significant delta except Noga^249^ (Fig 7b). Similar to the calculated amplitude (based on NPQlss rhythmicity) the physiological traits of Chl F are significantly influenced by the plasmotype diversity both under OT (P=0.0002 and P<0.0001) and HT (P<0.0001) (Fig S3c and d) and all cytolines show high thermal plasticity (Fig 7c and d).

**Figure 7:**
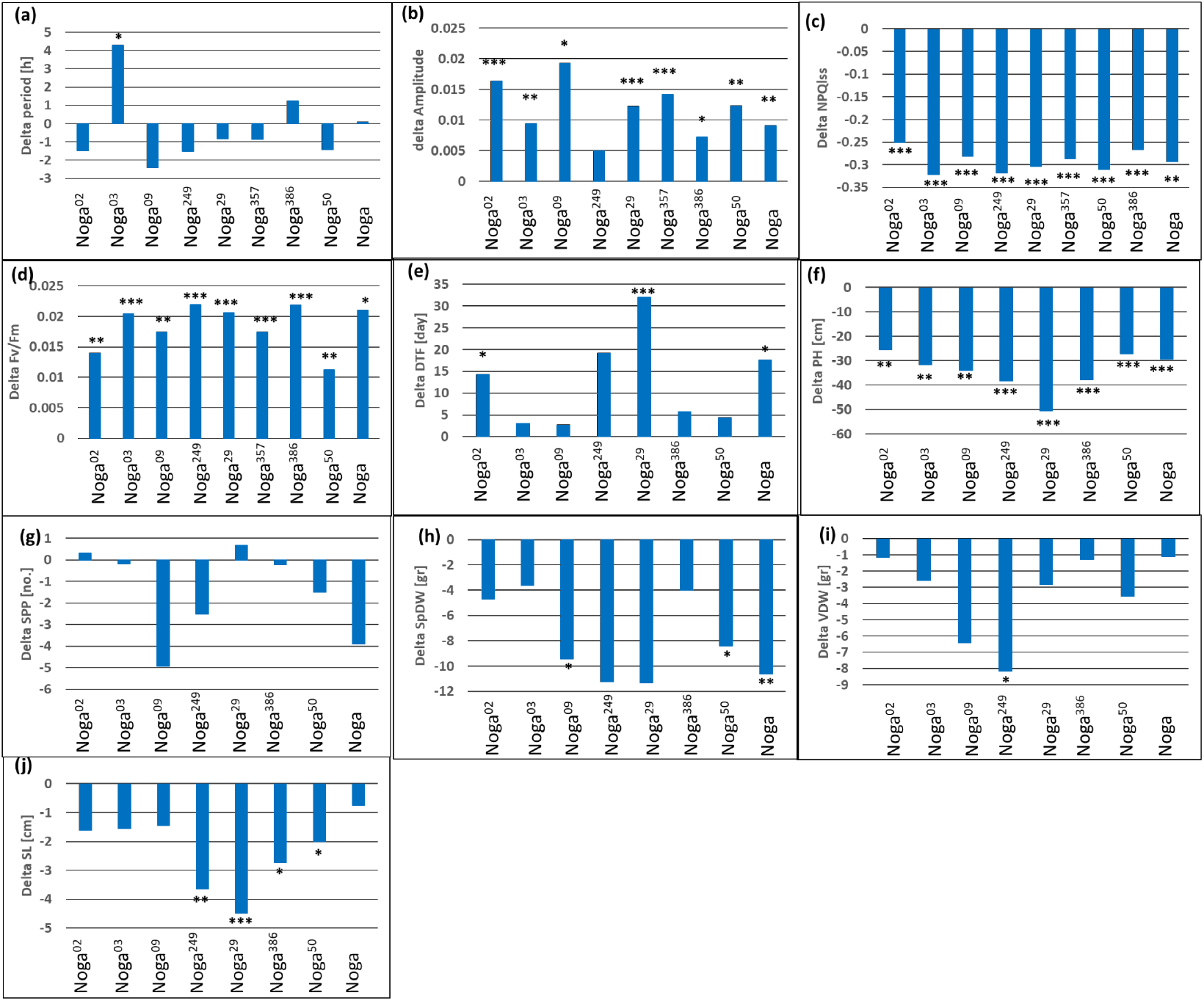
Phenotypic variation and thermal plasticity of cytolines for clock rhythmicity, Chl F, and life history traits. Bar plots for the delta between high temperature (HT, 32°C) and optimal temperature (OT, 22°C) (HT-OT/AT) for cytolines with wild barley plasmotype in the background of cultivated barley phenotype. For clock traits in the SensyPAM under HT and OT: **a)** period, **b)**, amplitude, **b)** NPQlss, and **d)** Fv/Fm. For life history traits under AT and HT: **e)** days to flowering (DTF), **f)** plant height (PH), **g)** spikes per plant (SPP), **h)** spikes dry weight (SpDW), **i)** vegetative dry weight (VDW), **j)** and spike length (SL). The Students’s t test p value is depicted as *, P<0.05; **, P<0.01; or ***, P<0.001.

We tested cytolines also in nethouse under AT and HT conditions. Differences between cytolines for DTF were significant only under HT (average of 117 ±17.4 days) including a delay in flowering and no difference was detected under AT (average of 107 ±7.8 days). The difference in DTF between cytolinesis significant under AT (P<0.0001), but no significant difference under HT, presumably since less plants were involved and not all the cytolines reached the flowering stage due to extreme temperatures observed during the year 2021 (Fig. S1). Plants were significantly higher under AT (PH=105.6 ±10.4 cm) compared to HT (72.2 ±12.6 cm) with all cytolines showing a significant plasticity (Fig 7f). Nevertheless, in both AT and HT treatments the cytolines were not significantly different for PH, SPP, SpDW, VDW and SL (Tukey-Kramer test; P<0.05) (Fig S4b to f). There was no significant difference for Spikes per plant (SPP) between AT (13.64 ±5.4) and HT (12.1 ±5.9) and no significant delta (Fig. 7g). Cytolines Noga^09^ and Noga^249^ are more similar to Noga as compared to delta small difference indeed observed among other lines between treatments. There was a significant reduction in the SpDW between AT (16.7 ±7.5gr) and HT (9 ±6.3 gr) (Fig S4d), depicting the heat’s clear detrimental effects on reproductive fitness. There is a large difference in SpDW-related delta values between the three cytolines Noga^02^, Noga^03^, and Noga^386^, andNoga (Fig 7h). VDW is significantly different between AT (21.16 gr) and HT (17.8 gr) and the delta values for Noga^09^ and Noga^249^ differ a lot from the values observed for Noga. VDW shows a highly significant decrease between AT (21.2 ±8.2 gr) and HT (18.1 ±7.1gr) and high variance exists between cytolines regarding their delta values. However, only Noga^249^ differs in delta from Noga. SL also changes significantly between AT (11.2 ±1.3 cm) and HT (9.1 ±1.9 cm), and there is high variance between the cytolines, Noga^249^, Noga^29^, Noga^386^ and Noga^50^ cytolines being significantly different in delta from Noga (Fig. 7j).

### Candidate chloroplast diversity underlying traits variation

In our previous study (Bdolach et al., 2019) we obtained and compared the chloroplast sequences of B1K-09-07 and B1K-50-04, which represent the parental lines of the ASHER doubled haploid population. Here, we expanded the collection of wild barley chloroplast sequences and included nine additional accessions (see Methods), in an attempt to associate the variation in clock and life history traits observed in the diallel to organelle genome diversity. Since our diallel doesn’t meet the population size criteria necessary for GWAS, we performed a Student’s t-Test for each clp haplotype with the different traits and corrected for the multiple testing (number of haplotypes; see Methods). Sequence alignments of the 11 chloroplast genomes identified 11 distinct haplotypes which include one to three genes (Table S2). Overall, we could observe that among the diallel hybrids, clp haplotypes are more significantly associated with variation in reproductive, compared to other trait types. (Fig 8). Previously, the comparison between Ashkelon and Hermon’s chloroplast genomes (B1K-09-07 and B1K-50-04) identified a non-synonymous SNP at the *rpoC1* gene (position: 24553; N571K) and we speculate that this gene could be responsible for the clock difference between the two subpopulations within ASHER (Bdolach et al., 2019). In the current diallel, the *rpoC1* and *matK* (position: 2099) co-segregate and this *matK/rpoC1* haplotype is significantly associated with DTF under AT and HT (P <0.0001) but not with the clock traits. However, another member of the PEP complex (Hess et al., 1993; Gajecka et al., 2021), i.e. *rpoC2*, appeared as a significant QTLs for several growth and reproductive traits. Within *rpoC2* we identified four SNPs (positions: 26445, 26808, 28702 and 29415) with the first two SNPs being in full linkage disequilibrium (LD), i.e. they are co-segregating between lines. The third and fourth *rpoC2* SNPs are each in LD as well, either with *ndhC* (position: 49896) or with *atpl* (31364) and *rps3* (80078). Within the diallel, the first *rpoC2* haplotype (namely *rpoC2*) is significantly associated with ASDW under both AT and HT for (p<0.0003 and 0.003, respectively), DTF (p<0.0007 and 0.0012, respectively), SL (0.0009 and 0.0004, respectively) and VDW (0.0014 and 0.0004, respectively) and only under AT it is associated with TDM (p<0.0012). The second haplotype, *rpoC2/ndhC*, is only significant for SL under AT (0.0004). The *rpoC2/atpl/rps3* haplotype is also significant under both AT and HT for SpDW (p<0.0001 and 0.0022), TDM (p<0.0001 and 0.0015) and VDW (p<0.0017 and 0.0049). This *rpoC2/atpl/rps3* haplotype is significant (p<0.005) for the clock trait dAmplitude. In *atpB* we identified two SNPs (positions: 52210 and 52297). For *cemA* and *ndhF*, and for *petB*, there was no significant association to any of the phenotypes. We identified *infA* and *ndhD* haplotypes as a significant QTLs (p<0.0053) for NPQlss under HT. To summarize, in this diallel analysis we identified more diversity linked with reproductive traits, e.g DTF, than for clock traits. Nevertheless, significant association between the rpoB/rpoC2/atp haplotype and clock amplitude plasticity (delta Amp) could be observed.

**Figure 8:**
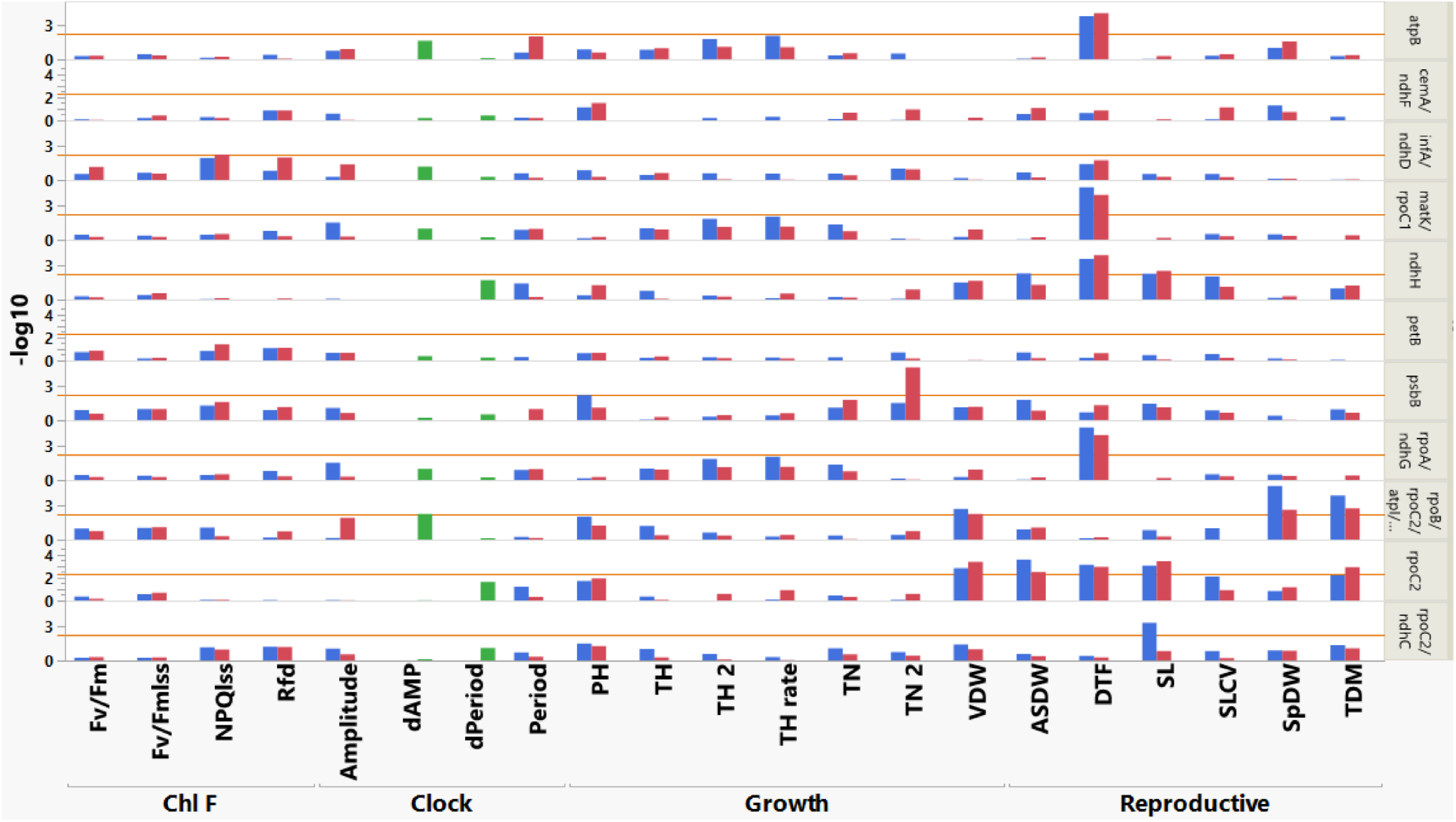
Genetic association between chloroplast haplotypes (right) and different clock and fitness traits (X-axis). Blue and red bars represent the -Log_10_P for Students’s t test between haplogroups among the hybrids under OT/AT and HT, respectively. The -LogP is calculated and corrected for multiple testing and the threshold for P<0.05 (-Log10=2.3) is indicated with horizontal orange line.

## DISCUSSION

### Whole plant and circadian clock responses to high temperatures and their interrelationship vs pleiotropy

In this study we found significant responses of plant growth and clock rhythmicity under elevated temperature. Comparison between early and late growth phenotypes showed that plants growing in high temperatures initially gain some growth advantage, as reflected by the higher tiller heights measured at about one month after transplanting. However, at the final time of harvest the heat is correlated significantly with reduced height, biomass and reproductive output (Fig. 1). Also, we found that heat is related with loss of robustness of the growth as could be viewed in the significant elevated CV of the spikes. Interestingly, the population of inbreds (ASHER DH) seemed to be more affected than the diallel hybrids, with the latter maintaining, for example, a similar mean for PH and SL values. These differences in the stability of hybrids was reported in many plant species and might be related with higher allelic heterogeneity across the genome (Fridman, 2015), which to some extent may allow the plant to show a wider reaction norm as suggested in biochemical models of heterosis (Goff, 2011).

Regarding the circadian clock output, which is measured at the early stages of development during the transition to flowering (leaf 3-4), the heat exerted a significant effect in both populations (ASHER (Bdolach et al., 2019) and diallel; Fig. 3), whereby the trends observed in the diallel are similar to those reported before for the DH population, with a mean acceleration of the clock by 1.97 hr and reduction of the amplitude by 2.4%. Although all traits were affected by the heat, the correlation matrix between them did not find many significant correlations between plasticity of clock and growth traits across either DH lines or hybrids. The link, however, could be established by pleiotropy at several loci such as *frp2.1* or *amp7.1* in the ASHER population, and with those QTLs CNI on the clock as well as vegetative and reproductive output of the plants (Fig. 6). Genetic correlations among traits arise from the pleiotropic effects of genes on multiple traits and/or linkage disequilibrium among distinct loci, each affecting a single member of the character complex (Flaconer and Mackay, 1996). The major differences between lack of relationship by genetic correlation (Table S1) to one found by pleiotropy (Fig. 6) are probably the additional genetic correlations with unmeasured traits (Gellman and Turner, 2020), or de facto inclusion of additional causal loci on either of the traits. For example, Vishnukiran et al. (2020) report major pleiotropic QTLs in rice between straw nitrogen and yield while there was no correlation between these two complex traits.

### Nature of the plasmotype natural diversity contribute to phenotypic variance

The genetic association we performed between the DNA diversity found among the wild barley in the ASHER or diallel population point to a significant effect of several haplotypes on the pleiotropic effects of clock and life history traits (Fig. 6 and Fig. S2). Previously, we reported on a non-synonymous variation in *rpoC1* (N571K) as a possible source for the significant differences in the clock plasticity between carriers of the plasmotype of B1K-50-04 and B1K-09-07 (Bdolach et. al. 2009). The plastidial *rpoC1* protein is a subunit of the holo-PEP complex (plastid encoded polymerase) known to interact with sigma factor 1-6, out of which at least SIG5 was shown to regulate rhythms of gene transcription, e.g., psbD (Noordally et al., 2013). Moreover, the PEP complex includes additional proteins encoded by chloroplast genes (Pfannschmidt et al., 2015) for which we identified association with pleiotropic effects on life history and clock traits (Fig. 8). This includes the link between diversity at the *rpoC2* and *rpoB* genes with the amplitude variation, mostly under HT.

Zooming in on this significant and hitherto unknown relationship between PEP variation and clock thermal plasticity will require a more thorough analysis of more advanced and isogenic lines. In the PEP complex, one major functional group is comprised of PAPs involved in DNA/RNA metabolism and gene expression regulation, while the second group is related to redox regulation and reactive oxygen species protection (Steiner et al., 2011). Moreover, the PEP is somehow coordinated with the nuclear encoding RNA polymerase (Pfannschmidt et al., 2015). Therefore, presumably non-synonymous variations (such as those between *rpoC2* alleles in current study) could be as effective as non-synonymous ones (between *rpoC1* alleles) in the functionality and variation we observed. It would be therefore required to look at different layers (transcriptome, proteome) between nearly isogenic and not necessarily knockout mutant lines to achieve relevant causal variation. Recent developments in plastid gene editing, also in cereals, may assist in generating and analyzing both types of mutations in barley and learn how they might modulate physiology and development of the plant under normal and high temperatures. Recent experiments suggest that most recent developments of TALLEN-based allele editing tested in *Arabidopsis* (Nakazato et al., 2021) could also be applied in barley (Fridman and Arimura, Personal communication) to allow such multi-layer analysis of isogenic mutants.

### Candidate genes in the frp2.1 and amp7.1 loci

We identified 48 and 71 high confidence genes in *frp2.1* and *amp7.1*, respectively. In the Barley NET we identified 751 genes that are interacting with the core clock genes in barley with scores ranging between 16 (highest) to 1.12 (lowest; Table S3-6). Within the *amp7.1* QTL region, we found four candidate genes including HORVU7Hr1G083270 (WRKY DNA-binding protein 70, score 1.89), HORVU7Hr1G083360 (NAD-dependent epimerase/dehydratase, score 1.42), HORVU7Hr1G084240 (transcription factor HY5, score 1.31), and HORVU7Hr1G084310 (overexpressor of cationic peroxidase 3, score 3.27). The guide (core circadian) genes for these interacting genes are PRR95 and PRR7 In *Arabidopsis*, the *HY5* binds with the G-box element of the *Lhcb* promoters thus indicating that *CCA1* can alter HY5-binding to the G-box through a direct protein–protein interaction in *Lncb* and *CCA1* (Andronis et al., 2008). Furthermore, the absence of *HY5* leads to a shorter period of *Lhcb1*. This suggest that interaction of the *HY5* and *CCA1* proteins on *Lhcb* promoters is necessary for normal circadian expression of the *Lhcb* genes, which may be related to the F-based measurements in current study. Regarding the *frp2.1* QTL region, we found only one candidate interactive locus i.e. HORVU2Hr1G103620 (ABC transporter C family member 2, Score-1.87). Notably, mining the allelic diversity of these candidate genes within the larger Barley1K GWAS panel (Hubner et al., 2009) provides further support to their role in the manifestation of the rhythmicity of the clock output, and its plasticity under high temperature. For example, the *frp2.1* region was also found in association with period under HT in the larger B1K panel (Manuscript in preparation).

It may well be that implementation of two-dimensional QTL studies in larger populations will validate the observed cytonuclear interactions (Fig. 6; Fig. S2) however, it will require a larger scale of Barley1K chloroplast sequencing. These *in silico* identified interactions between candidate loci can then be further verified in *in-vivo* interaction studies that would expand our knowledge of the circadian clock network and its role in heat sensing and plant responses.

### The potential of ancestor plasmotype and cytonuclear diversity for crop improvement

The potential of plasmotype diversity for breeding better adapted barley could be considered from the perspective of several important traits relevant to adaptation to different environments. Based on the reciprocal hybrids and cytolines, our results clearly show that flowering time is perhaps the trait most affected by plasmotype diversity. For example, the flowering of cytoline Noga^03^ under HT is not significantly delayed as compared to the cultivated Noga reference, which is flowering more than two weeks later under the same conditions (Fig. 7e). This robustness includes reduced effects of the heat on the reproductive output (Fig. 7h). While these plasmotype alleles bear the potential to increase crop fitness and broaden the environment in which we can grow a major crop plant, the cytonuclear interactions are as important to consider. The mutual conditioned effects of nucleotype and plasmotype QTL (Fig. 6) indicate that a more extensive genetic infrastructure is required to capture both types of wild alleles in a cultivated genetic background in order to allow field-testing. The current multi-parent populations in cereals and barley do not include cytonuclear interactions segregation (Schnaithmann et al., 2014; Maurer et al., 2015; Novakazi et al., 2020). Therefore, we recently developed a barley interspecific cytonuclear multi-parent population (CMPP) with the goal of studying CNI and its utility for breeding and for testing pleiotropic effects on clock rhythmicity and thermal plasticity.

## CONCLUSIONS

The ability to test clock phenotypes on the same populations that grow in the field and identify the underlying genetics is key to understanding the relationships between important plant traits and circadian clock mechanisms. (here, and reviewed in Panter et al., 2019). The low occurrence of significant correlations between clock and fitness traits yet the existence of significant pleiotropic QTLs (Prusty et al., 2021) highlight the complex nature of circadian clock rhythmicity and yield traits. Several studies show the effect of clock gene mutants on crop behavior in natural and agricultural environments (Izawa et al., 2011; Bendix et al., 2015). Here, we show that accounting for plasmotype diversity, which modulates the plasticity of clock output, has the potential to confer yield robustness under adverse thermal conditions Pinpointing the underlying pleiotropic genes is a key to further unravelling the interplay between core and output clock pathways, which may work in both directions through mechanisms yet to be discovered.

## MATERIALS AND METHODS

### Plant material

The source for the ASHER and diallel populations described in this study are barley accessions (*Hordeum vulgare* ssp. *spontaneum*) that we selected from the Barley1K collection in Israel to represent the different genetic clades (Hubner et al., 2009). In addition, few lines are from the IPK collection (Maurer et al., 2015). Included also in the diallel and as cultivated background for making cytolines is the *H. vulgare* cultivar cv. Noga which is the leading barley line in Israel. The wild accessions are from Yerucham (B1K-02-02), Michmoret (B1K-03-09), Ein Prat (B1K-04-04), Neomi (B1K-05-07), Ashqelon (B1K-09-07), Mount Arbel (B1K-29-13), Mount Harif (B1K-33-09), Jordan Canal (B1K-42-16), Mount Eitan (B1K-49-19), Mount Hermon (B1K-50-04), Kisalon, Israel (HID386), Turkey (HID357) and Iran (HID249). The ASHER is an F3 DH population generated from two reciprocal hybrids between Ashkelon (B1K-09-07) and Mount Hermon (B1K-50-04) and we described it in detail earlier (Bdolach et al., 2019b). The diallel is a reciprocal cross scheme between 11 wild accessions (from the B1K and HID386) and Noga that were intercrossed on each other to create a full set of hybrid pairs that differ in their plasmotypes. To generate the cytolines in the background of cultivar Noga we continued with the F1 hybrids that carry the wild plasmotypes and kept backcrossing it to Noga as male for several generations. As an example, Noga^02^ is a cytoline that carry the plasmotype of B1K-02-02 after we performed five backcrosses followed by three generations of selfing to achieve BC_5_S_3_ line. More cytolines are Noga^03^, BC_5_S_2_ of Noga x B1K-03-09 cross; Noga^09^, BC_5_S_2_ of Noga x B1K-09-07 cross; Noga^249^, BC4S2 of Noga x HID249 cross; Noga^29^, BC5S3 of Noga x B1K-29-13 cross; Noga^386^, BC_3_S_2_ of Noga x HID386 cross, and Noga^50^, BC_3_S_2_ of Noga x B1K-50-04 cross.

### Growth and phenotyping

We conducted the net house experiments in the Agricultural Research Organization - Volcani (ARO) Center, Israel. We sowed the different lines in germination trays and at the 3-leaf stage transplanted the seedlings in randomized block design into troughs measuring 0.4 × 0.3 m (Mapal Horticulture Trough System, Merom Golan, Israel). A trough contained two rows of plants and the soil was composed of two layers of volcanic soil (4–20 type of rough soil topped by a finer Odem193 type; Toof Merom Golan, Merom Golan, Israel). We applied irrigation and fertilization using a drip system (2L per hour, every 30 cm) four times a day for 10 minutes. Due to the sensitivity of wild barley to day-length conditions, we preferred to achieve mild higher temperature conditions by warming the nethouse rather than late sowing conducted for example for tomato (Bineau et al., 2021). We achieved high temperature treatment (HT) by covering half of the insect-proof with nylons and heating with electric heathers (3KW; Galon fans and pumps Ltd, Nehora, Israel). The second half of the nethouse remained with only net walls and ventilated with a large fan to take out the hot air for the ambient temperature treatment (AT). The thermal differences between HT and AT is depicted in fig. S1, with a mean increase of 3.9 °C and 2.8 °C during day and night time and maximum delta of mean 7.5 °C between AT to HT.

We measured circadian clock amplitude and period in high-throughput SensyPAM (SensyTIV, Aviel, Israel) custom-designed to allow Fluorescence measurements (Bdolach et al., 2019) under optimal temperature of 22°C (OT) or high temperature of 32°C (HT). We calculated the Fluorescence parameters NPQlss, Fv/Fm, Fv/Fmlss and Rfd as average of all three days measurements under continuous light (Dakhiya et al., 2017). We calculated the period and amplitude of the circadian clock output using the BioDare platform (https://biodare2.ed.ac.uk) (Zielinski et al., 2014).

We obtained the life history traits phenotype for the ASHER population lines during winter of 2017-2018 in six replicates per treatment. The reciprocal diallel population were grown during winter of 2019-2020 and the cytolines experiment was conducted in winter of 2020-2021. We began phenotyping by measuring Tiller height (TH), that is the length of the longest tiller from ground level to the last fully expended leaf in that tiller. Tiller number (TN) is the number of tillers per plant and it was determined about one month after transplanting the plants. TH and TN ware measured once (_1) or twice (_2) with 14 days apart. We calculated TH rate by suspecting TH_2 with TH_1 and dividing with the number of days between these two measurements. We determined the number of days to flowering (DTF) based on the date when the first awns appear in the main tiller. During grain filling we measured five spikes per plant for spike length (SL) and later to obtain SLCV. In addition, during grain filling we measured plant height (PH) from ground to the start of the toolset spike. We then cached the five and whole spikes of each plant in separate paper and nylon bags, respectively. Plants ware left to dry for several weeks after irrigation was terminated. We harvested dry plants by cutting at soil level and placing them in the nylon bags. Weight of the nylon bag with the plant is the total dry matter (TDM). We collected dispersal units from bag and weighted them. We calculated average spike dry weight (ASDW) based on weighing the five spikes that we cached in the paper bag. We then summed the weight of spikes (dispersal units) in the plastic and paper bags to obtain spikes dry weight (SpDW). Vegetative dry weight (VDW) is the reduction of SpDW from TDM. In the cytolines experiment, we also counted Spikes per plant (SPP) and the ASDW based on those spikes.

### Genome-wide and cytonuclear interaction QTL analysis

The description of the ASHER SNP genotyping and QTL analysis for the different traits is described in Bdolach et al., 2019. The genome-wide QTL interaction analysis of the DH population for different traits carried out using inclusive composite interval mapping (ICIM; (Li et al., 2007)) with the IciMapping V4.1 (Meng et al., 2015) software package. IciM 4.1 uses an improved algorithm of composite interval mapping for bi-parental population. The QTL by environment interaction (QxE) was also assessed with the inclusive composite interval mapping (ICIM) method, using the MET function of the software QTL IciMapping 4.1 (Li et al., 2007, Meng et al., 2015). Illumina paired-end libraries (375 bp insert size) of total barley DNA from mature leaves were used to sequence the plastid genomes of the parental lines as previously described (Bdolach et al., 2019).

### Statistical analysis

The JMP version 14.0 statistical package (SAS Institute, Cary, NC, USA) was used for statistical analyses. Student’s t-Tests between treatments, plasmotypes and alleles were conducted using the ‘Fit Y by X’ function. A factorial model was employed for the analysis of variance (ANOVA, Fig. 6), using ‘Fit model’, with temperature treatment and allelic state as fixed effects.

### Candidate genes in the frp 2.1 and amp7.1

We downloaded the list of high confidence gene in the QTL intervals (*frp2.1*, Chromosome 2:698,875,542-702,308,910; *amp7.1*, Chromosome 7: 498,472,330-510,903,725) from BarleX database with Morex V1 annotation and tested them for interaction with core clock gene in barley. For this, we retrieved the list of genes involved in circadian pathway in *Hordeum vulgare* from plant reactome (https://plantreactome.gramene.org). Plant reactome is the Gramene’s pathway knowledgebase that uses *Oryza sativa* as a reference species for manual curation of the pathway and extends pathway knowledge for other 82 plant species via gene-orthology projection (Naithani et al., 2020). BarleyNET inferred the co-functional links between barley genes by analyzing various types of omics data obtained from cultivated barley, as well as three other plant species (*Arabidopsis thaliana, Zea mays*, and *Oryza sativ*a) (Lee et al., 2020). In the BarleyNET, under the pathway centric search function, the known circadian clock genes were used as the guide gene to identify genes by ‘guilt-by-association’ method. These genes were prioritized by total edge weight score (sum of log likelihood score) to the guide gene set.

## ACKNOWLEDGEMENTS

We thank Dr Stephan Greiner (Max-Planck-Institut für Molekulare Pflanzenphysiologie, Golm, Germany) for sharing barley chloroplasts sequence data. The authors are grateful to Royi Levav Oded Anner and Daniel Shamir (SensyTIV, Amiel, Israel) for their assistance in maintaining the SensyPAM as a system for measuring circadian rhythms. We also wish to thanks the technical assistance of laboratory member Avital Beery and Orit Amir-Segev.

## LIST OF AUTHOR CONTRIBUTIONS

E.B and E.F. designed the experiments, collected, analyzed and interpreted data, and wrote the manuscript. E.B., M.R.P. K.K., and L.D.T were involved in the data analyses, their interpretation and in writing the manuscript.

---

**Table S1:** Pearson correlations (r) between all phenotypic traits under optimal temperature (OT, 22°C) and high temperature (HT, 32°C) in the reciprocal dialle population.

Table S2: Chloroplast genes sequencing of the *H. spontaneum* accessions parental accessions of the diallel: B1K-02-02 (Yerucham), B1K-03-09 (Michmoret), B1K-04-04 (Ein Prat), B1K-05-07 (Neomi), B1K-09-07 (Ashqelon), B1K-29-13 (Mount Arbel), B1K-33-09 (Mount Harif), B1K-42-16 (Jordan Canal), B1K-49-19 (Mount Eitan), B1K-50-04 (Mount Hermon), HID386 (Kisalon, Israel) and the cultivar Noga.

**Table S3:** Barley circadian pathway genes from the plant reactome (https://plantreactome.gramene.org)

**Table S4:** List of the interacting genes to the guide core circadian genes from BarleyNET pathway centric search

**Table S5:** High confidence candidate genes in *amp7.1* chromosomal region

**Table S6:** High confidence candidate genes in *frp2.1* chromosomal region

**Figure S1**: Mean daily temperature in the nethouse under ambient temperature (AT), high temperature (HT) and the delta between them (HT-AT). **a)** ASHER, **b)** diallel and c) cyolines experiments. Blue and orange lines represent the average day of flowering under AT and HT, respectively.

**Figure S2: GxGxE interactions between *amp7.1* and plasmotype under optimal and ambient temperatures.** Cytoplasm by nuclear (GxG) QTL interaction plots under **a)** ambient (AT) and **b)** high (HT) temperatures for Days to flowering. Plasmotype is depicted in X axis (Ashkelon or Hermon), and red or blue lines illustrate the Hermon or Ashkelon alleles in *amp7.1*. Levels not connected by same letter are significantly different in student’s t-test. **c)** Reaction norms of the different combinations Plasmotype-*amp7.1* loci between optimal temperature (OT, 22°C) and high temperature (HT, 32°C). Green, Ashkelon (Plasmotype)-Hermon (*frp2.1*); Purple, Ashkelon-Ashkelon; Brown, Hermon-Ashkelon; Ornage, Hermon-Hermon

**Figure S3: Clock rhythmicity and Chl F variation between cytolines under OT and HT**. Cytolines with wild barley cytoplasm in the background of cultivated barley grown under optimal temperature (OT, 22°C) and high temperature (HT, 32°C). Bar plots with SE for the cytolines for **a)** period, **b)** amplitude, **c)** NPQlss and **d)** Fv/Fm. Different letters depict significant difference in a Tukey-Kramer test.

**Figure S4: No different between cytolines for life history traits under AT and HT**. Cytolines with wild barley cytoplasm in the background of cultivated barley grown under ambient temperature (AT) and high temperature (HT). Bar plots and SE for **a)** days to flowering, **b)** plant height, **c)** spikes per plant, **d)** spikes dry weight, **e)** vegetative dry weight and **f)** spike length. Different letters depict significant difference in Tukey-Kramer test.

## Notes

This work was supported by grants from the Israel Science Foundation (ISF 444/21) and Horizon2020/CAPITALISE AMD-862201-2 grants to E.F.

